# Human hippocampal ripples predict the alignment of experience to a grid-like schema

**DOI:** 10.1101/2025.01.08.632069

**Authors:** Zhibing Xiao, Xiongfei Wang, Jinbo Zhang, Jianxin Ou, Li He, Yukun Qu, Xiangyu Hu, Tim Behrens, Yunzhe Liu

**Affiliations:** State Key Laboratory of Cognitive Neuroscience and Learning, IDG/McGovern Institute for Brain Research, Beijing Normal University, Beijing, China; Chinese Institute for Brain Research, Beijing, China; Department of Neurosurgery, Laboratory for Clinical Medicine, Sanbo Brain Hospital, Capital Medical University, Beijing, China; Wellcome Centre for Human Neuroimaging, UCL, London, UK; Wellcome Centre for Integrative Neuroimaging, University of Oxford, Oxford, UK; Sainsbury Wellcome Centre for Neural Circuits and Behaviour, UCL, London, UK

**Keywords:** hippocampal ripples, cognitive map, grid code, offline learning, DMN

## Abstract

Humans create internal cognitive maps that allow us to make inferences beyond direct experience. These maps often rely on hexagonal grid-cell-like neural codes, serving as a schema for two-dimensional (2D) spaces. Yet it remains unclear how new experiences become aligned with this schema, especially in non-spatial contexts. Here, we show that hippocampal ripples – brief bursts of neuronal activity during rest – predict the emergence of grid-like codes in a novel 2D inference task. We recorded intracranial neuronal activity in 42 epilepsy patients as they learned rank relationships among feature objects (for example, objects differing in “magic” or “speed”). After learning, these objects were combined to form “compounds” occupying a 2D conceptual space defined by two feature dimensions. During learning, hippocampal ripple activity increased during pauses between trials, suggesting that ripples integrated newly acquired information offline. Subsequently, ripple activity during post-learning rest predicted the later emergence of grid-like codes in the entorhinal cortex (EC) and medial prefrontal cortex (mPFC), a core region of the default mode network (DMN), when participants inferred unseen relationships among the compounds. Critically, coordination during rest between hippocampal ripples and DMN activity in the mPFC predicted participants’ ability to infer complex relationships beyond direct memory retrieval. These findings provide the first direct evidence that hippocampal ripples, working with the DMN, align new experiences with a grid-like schema offline, transforming discrete learning events into structured knowledge that supports flexible and adaptive reasoning in human cognition.

## Introduction

Imagine reading a complex novel filled with many interconnected characters. As you learn about their individual relationships, you begin to form a mental map that allows you to infer connections not directly described. This ability to build structured knowledge and make inferences beyond direct observation is a hallmark of human intelligence.

Central to this capacity are cognitive maps: internal representations of how different concepts relate to one another. In animals, researchers have identified grid cells in the brain that represent a neural map of physical space, characterised by a distinctive six-fold, hexagonal pattern (Hafting et al., 2005). In humans, similar grid-like codes have been detected not only for physical navigation (Doeller et al., 2010; Jacobs et al., 2013) but also in abstract, non-spatial domains, located in regions such as the entorhinal cortex (EC) and medial prefrontal cortex (mPFC) (Constantinescu et al., 2016; S. A. Park et al., 2021; Qu et al., 2024). These grid-like codes are thought to function as a neural coordinate system, helping us organise experiences and infer unseen relationships (Behrens et al., 2018).

The grid code is believed to serve as a stable scaffold for new experiences (Chandra et al., 2024; Whittington et al., 2020) and can be updated through hippocampal replay to align fresh inputs with existing representations (Bakermans et al., 2024). Understanding how such codes emerge in the human brain is crucial. However, no study has yet provided direct evidence of how grid codes are formed, especially for non-spatial spaces.

Capturing the emergence of grid-like representations is challenging because it requires observing neuronal activity from the very beginning of learning. A recent magnetoencephalography (MEG) study showed that neural replay – the sequential reactivation of experiences during rest – is essential for building cognitive maps (Ou et al., 2025). This replay is associated with power increases in ripple-band activity (Y. Liu et al., 2019, 2021). However, the limited spatial resolution of non-invasive neuroimaging reveals little about how these processes unfold within neuronal circuits or coordinate across different brain regions.

In rodents, replay events are often accompanied by hippocampal ripples – brief bursts of high-frequency oscillations during rest or sleep that are crucial for memory consolidation (Buzsáki, 2015; Buzsáki et al., 1992; Girardeau et al., 2009; Siapas & Wilson, 1998). Such ripples help form and refine spatial maps (Roux et al., 2017; van de Ven et al., 2016), integrate information to support future inferences (Barron et al., 2020), and coordinate with cortical areas including the EC (Ólafsdóttir et al., 2016) and mPFC (Kaefer et al., 2020). Whether a similar hippocampal–cortical mechanism underlies the alignment of new experiences with grid-like conceptual maps in humans remains unknown.

While hippocampal ripples in humans have mainly been studied in relation to episodic memory processes (Norman et al., 2019, 2021; Sakon et al., 2024) and, more recently, in spontaneous thoughts (Iwata et al., 2024; Kucyi et al., 2023), their role in learning, particularly in forming or aligning new experiences with grid-like codes in conceptual spaces, is largely unexplored. Demonstrating such a role would suggest that hippocampal ripples and grid codes form a core neuronal mechanism extending beyond physical navigation and across species, providing a scaffold for general task spaces.

In this study, we investigated whether hippocampal ripples during rest contribute to the alignment of new experiences with grid-like codes in humans. We worked with 42 patients who had intracranial depth electrodes implanted in both the hippocampus and widespread cortical regions, allowing us to record neuronal activity at high temporal and spatial precision.

We designed a novel task to track neuronal activity as participants learned to form the cognitive map of a two-dimensional (2D) conceptual space. Participants first learned the relative rankings of objects within each feature dimension, integrating information across adjacent items. During this learning phase, we observed that hippocampal ripples during inter-trial intervals (ITIs) increased over the learning experience, suggesting a role in integrating newly acquired information. After a 5-minute rest period, these feature objects were combined into compounds positioned within a 2D conceptual space defined by feature dimensions. Participants then made inferences about these compounds, requiring them to construct an internal representation, i.e., a cognitive map of the 2D space.

We found grid-cell-like codes in the EC and mPFC when participants made inferences about the 2D map, and hippocampal ripple activity during post-learning rest was directly linked to the emergence of these grid representations. Moreover, coordination between hippocampal ripples and mPFC activity during rest predicted inference abilities that went beyond direct memory retrieval. These results provide the first direct evidence that hippocampal ripples help align new experiences with grid-like conceptual schemas, transforming discrete learning events into structured knowledge for flexible and adaptive reasoning in human cognition.

## Results

### Task and intracranial recordings

To investigate the neural processes involved, especially during offline periods, in building a cognitive map, we worked with 42 patients who have refractory epilepsy, after excluding subjects who cannot perform the task, we have 38 patients (12 females, age: 25 ± 9.8 years, mean ± SD) in total for all subsequent analysis. These patients had intracranial electrodes implanted in the hippocampus and other cortical areas (e.g., DMN, see **Supplementary Tables 1, 2** for more details) as part of their neurosurgical evaluation.

Participants were first instructed to learn the hierarchical ranks within three feature dimensions: ‘magic’, ‘speed’, and ‘power’, each comprising four objects. The learning phase focused exclusively on adjacent pairs differing by one rank, referred to as ‘premise learning’. This phase is crucial, as two of the three learned features are later chosen to construct a two-dimensional (2D) task space (**Fig. 1A**). After the learning phase, subjects were tested on their memory performance regarding pairwise rank relationships and their ability to infer relationships spanning more than one rank, both within a feature dimension. Following a rest period, they underwent a re-test to assess both memory and inference abilities over the feature objects.

**Fig. 1.**
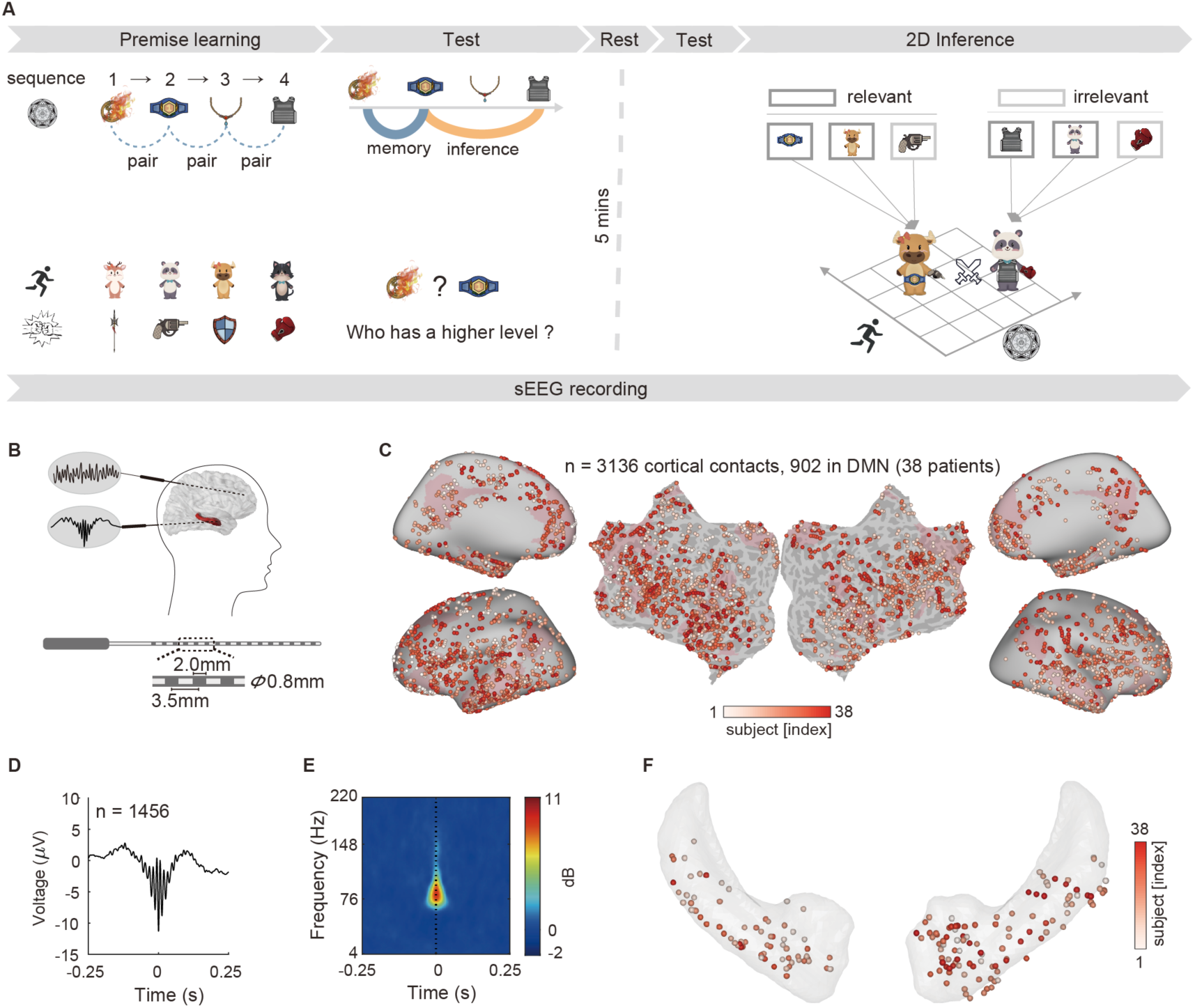
Task schematic, sEEG implantation and ripples. **A.** Subjects, undergoing stereo electroencephalography (sEEG) recordings, first learned the ranks of three feature dimensions and were later asked to perform inference on compounds composed of these feature objects. **B.** Schematic of intracranial electrode implantation for patients with epilepsy. **C.** Distribution of 3,136 contacts within 3mm of the cortical surface across 38 subjects, including 902 contacts within the Default Mode Network (DMN). Each contact is colour-coded based on the subject index. **D.** Waveform of hippocampal ripples observed in one demonstration contact. **E.** Time-frequency plot corresponding to the contact. **F.** Distribution of all 181 contacts within the hippocampus across subjects.

In the final phase, feature objects from the three feature sequences (one object from each sequence) were combined to form compounds. Unbeknownst to the subjects, these compounds were positioned within a 2D task space constructed upon two of the three learned feature dimensions (**Fig. 1A**). The specific relevant axes were randomised across subjects. Participants were asked to infer the unobserved relationships between compounds—a process referred to as *2D inference*—which requires an internal representation of the 2D task space.

Participants performed well on the feature test, with an average behavioural accuracy of 0.88 ± 0.12 (mean ± SD), which increased to 0.91 ± 0.10 after the rest period. For the 2D inference task, the mean accuracy was 0.86 ± 0.11.

We focused on the role of hippocampal–cortical communication in constructing the cognitive map. Using multi-contact intracranial electrodes, we recorded hippocampal ripples and cortical neural activity during the task and rest (see schematic in **Fig. 1B**). We analysed recordings from contacts within 3 mm of the cortical surface or the hippocampus, identifying 3,136 cortical contacts across 38 subjects, including 902 contacts within the DMN (**Fig. 1C**), and 181 hippocampal contacts within a subset of 34 subjects (**Fig. 1F**). Hippocampal ripples—transient high-frequency oscillations— were identified using established procedures (Norman et al., 2019; Stark et al., 2014; Vaz et al., 2019) (see Methods for details). An example of a hippocampal ripple waveform (**Fig. 1D**) and its corresponding time–frequency profile (**Fig. 1E**) are shown in **Fig. 1**.

### Hippocampal ripple activity increases as learning progresses

During premise learning, participants acquired knowledge about the hierarchical ranks of adjacent feature pairs. They were later tested on their ability to infer rank relationships that spanned beyond the directly learned pairs. Hippocampal ripples are hypothesised to integrate memories during offline periods, forming structures that support such inference (Barron et al., 2020).

To test this in the human brain, we analysed ripple rates at different phases of the trial: stimulus presentation, feedback, and the inter-trial interval (ITI) (**Fig. 2A**). The ITI represents an offline period during learning. The mean ripple rate during the ITI across all hippocampal contacts is shown in **Fig. 2B** (see also **Supplementary Fig. 1**). We found an overall higher ripple rate during the offline period (ITI) compared to the feedback periods (*Z* = 3.56, *P* = 3.65 × 10^-4^, linear mixed model, with the nested structure of subjects and contacts as random effect; see Methods for details).

**Fig. 2.**
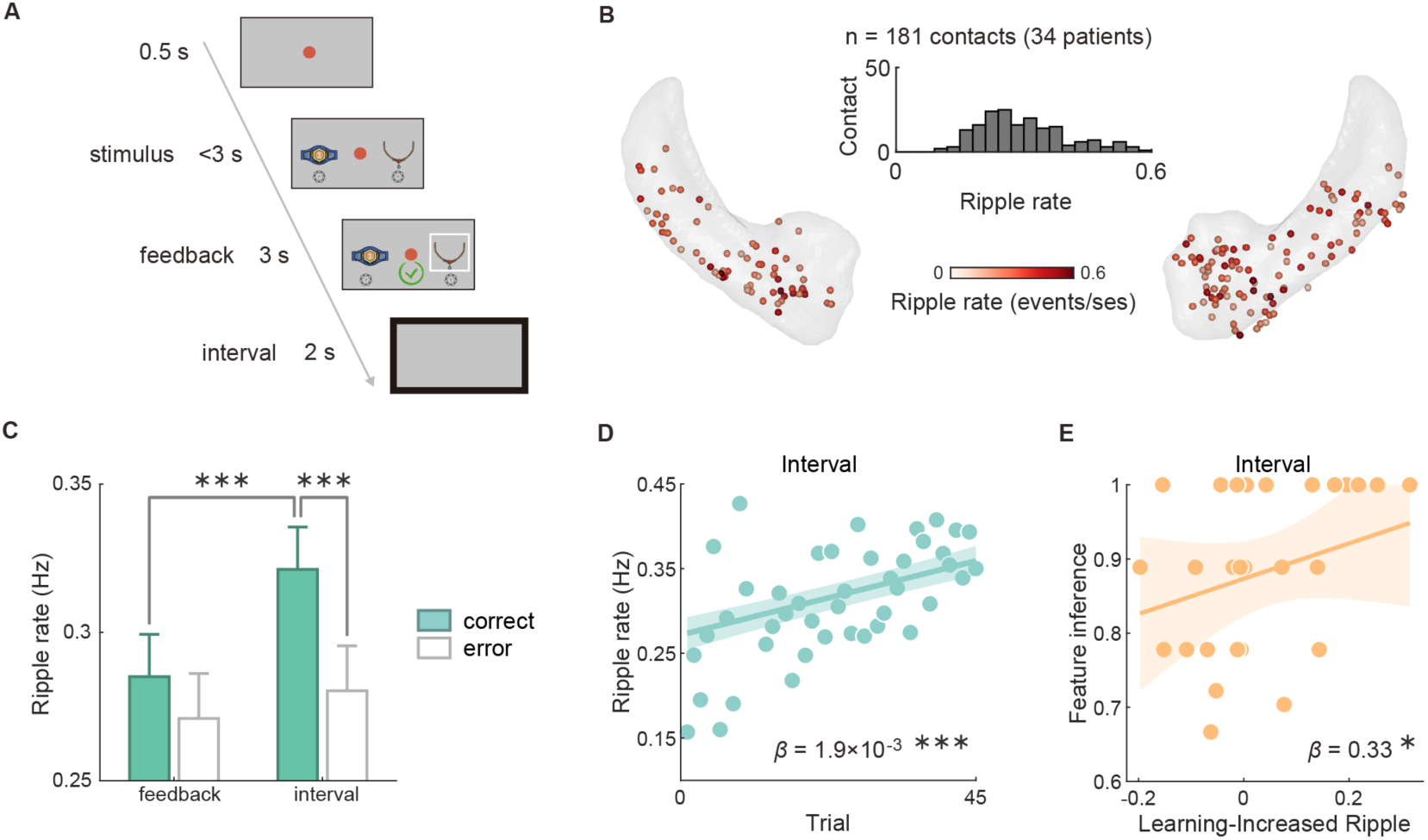
Hippocampal ripples facilitate offline learning. **A.** In an example trial of premise learning, participants were asked to select the relative higher rank from two alternative feature objects within 3-second time limit. Feedback was then presented with a correct/incorrect designation for 3-second, followed by a 2-second inter-trial interval (highlight in black rectangle). **B.** Ripple rates across all contacts during inter-trial intervals of learning in the hippocampus, with an inset plot showing their histogram. **C.** Ripple rates were higher during inter-trial intervals after correct responses than after error responses, and were also higher compared to those during feedback periods. **D.** Ripple rates during intervals following correct responses increased over the course of learning (number of learning trials). This increase in ripple rate as a function of learning is termed the Learning-Increased Ripple (Learn-R). Data analysis utilised a linear mixed-effects model with trial number as a regressor (see Methods for details). For illustration purposes, the mean ripple rate across subjects for each trial was presented in dots. See also **Supplementary Fig. 1**. **E.** Feature inference can be predicted specifically by the Learn-R (Learning-Increased Ripple) during the interval period. Each dot represents a subject. Error bars show SEM. * *P* < 0.05, *** *P* < 0.001.

We also found that ripple rates during the ITI were higher following correct responses compared to error trials (*Z* = 4.41, *P* = 2.05 × 10⁻⁵; Holm corrected; **Fig. 2C**), and this effect was more pronounced during the ITI than during the feedback period (*Z* = 4.80, *P* = 3.17 × 10⁻⁶; Holm corrected; **Fig. 2C**). No such outcome effect on ripple rates was found during stimulus presentation (**Supplementary Fig. 1D**). Notably, ripple rates during the ITI following correct responses increased over the course of learning (*β* = 1.9 × 10⁻³, *P* = 8.01 × 10⁻⁵; **Fig. 2D**), whereas no such increase was observed following error responses. We refer to this effect as the *Learning-Increased Ripple* (Learn-R). The Learn-R was calculated as the Spearman rank correlation coefficient (Fisher’s Z transformed) between the ripple rate during ITIs following correct responses and the trial number. We found that Learn-R was a unique predictor of feature inference performance, specifically during the ITI period (*β* = 0.33, *P* = 0.044; robust regression; **Fig. 2E**), but not during stimulus presentation or feedback periods. No relationship between Learn-R and memory performance was found across all learning phases (see also **Supplementary Fig. 1**). These findings suggest that hippocampal ripples during offline periods are involved in integrating experiences to support inference, highlighting the critical role of hippocampal activity during rest in learning processes.

### Increased hippocampal ripples during rest improve feature inference

After learning, participants took a 5-minute rest, followed by a re-test of their learning performance – memory (pairwise rank) and inference (more than one rank difference) both within a feature dimension (**Fig. 3A**). We hypothesised that the hippocampal ripple activity during this task-free period would further enhance inference performance.

**Fig. 3.**
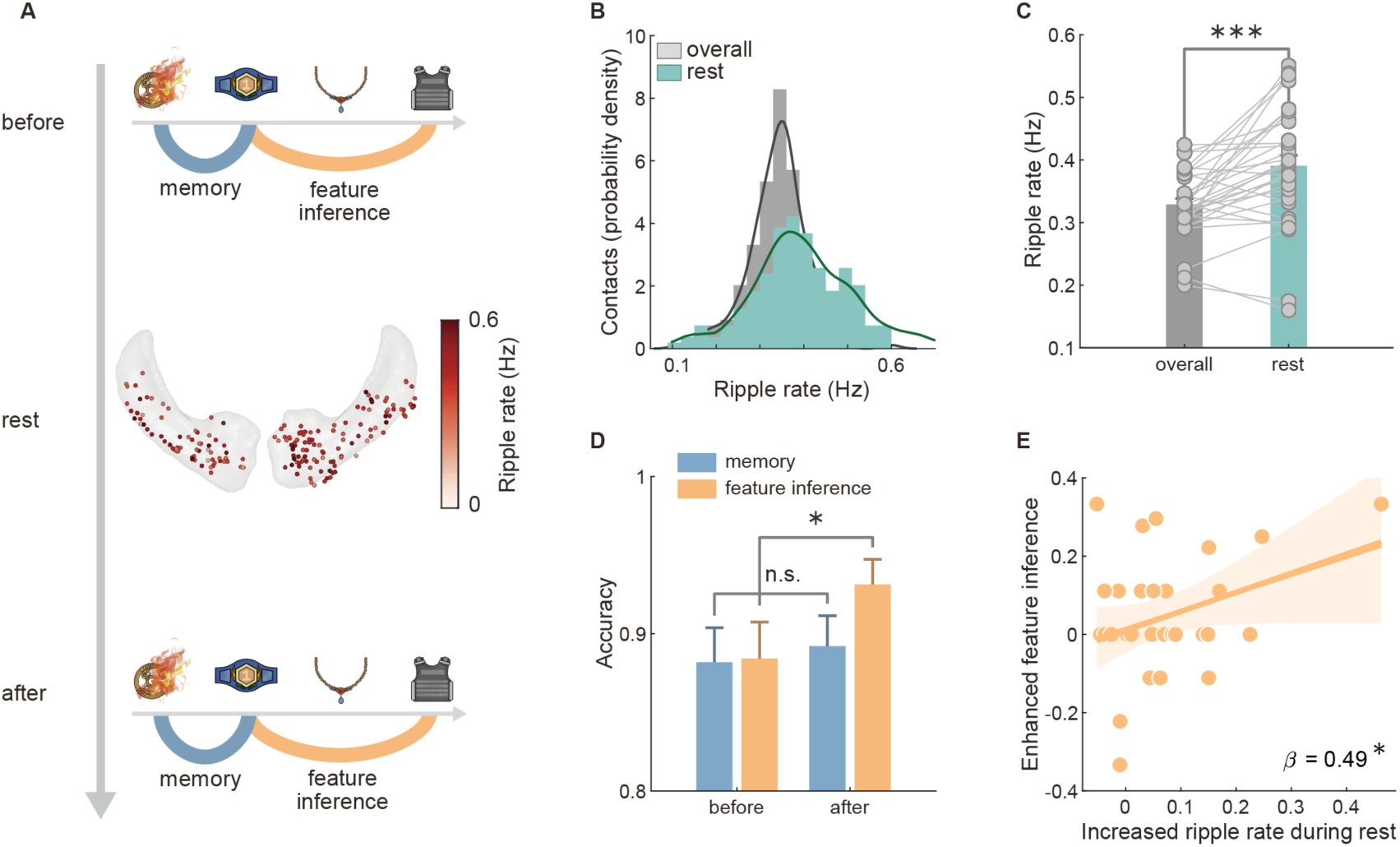
Hippocampal ripples during rest promote later feature inference. **A.** The same feature memory and inference tests were conducted before and after the rest period. During rest, the mean ripple rate of all contacts within hippocampus were shown. See also **Supplementary Fig. 2**. **B.** Distribution of ripple rates during rest compared to the mean ripple rates across the entire task. **C.** The mean ripple rates during rest were significantly higher than those observed throughout the entire task. **D.** After the rest period, the accuracy of the feature inference trials (non-adjacent pairs) increased significantly compared to the pre-rest period. This was not true for the memory tests (adjacent pairs). **E.** Positive correlation observed between the increased ripple rate during rest (compared to the overall mean) and the increased feature inference ability following the rest. Each dot represents a single subject. The solid yellow line indicates the linear fit. Error bars show SEM. * *P* < 0.05, *** *P* < 0.001, n.s., not significant.

We calculated the ripple rates during the 5-minute rest period for each contact (**Fig. 3B**) and then averaged them within subjects to analyse their association with behavioural performance. We found that ripple rates during rest were significantly elevated compared to the mean ripple rate across the entire task (*t*_(33)_ = 4.09; *P* = 2.58 × 10⁻⁴; **Fig. 3C**; see also **Supplementary Fig. 2**). After the rest period, participants’ feature inference performance improved significantly compared to before rest (*Z* = 1.78, *P* = 0.038; Wilcoxon signed-rank test; **Fig. 3D**).

Importantly, the improvement in feature inference performance was predicted by the increased ripple rate during rest (*β* = 0.49, *P* = 0.0419; robust regression; **Fig. 3E**). In contrast, memory performance did not change significantly after rest (*Z* = 0.62, *P* = 0.267; **Fig. 3D**), nor was there a correlation between changes in memory performance and increased ripple rate during rest (*P* = 0.214). These findings highlight the importance of hippocampal ripples during rest in facilitating the integration of learned information to enhance inference abilities.

### Ripples during rest predict later emergence of grid-like code on-task

Previous studies have demonstrated that human replay during rest facilitates the formation of cognitive map (Ou et al., 2025) and is associated with power increases in the ripple band (Y. Liu et al., 2019). In our study using iEEG, we investigated whether the identified hippocampal ripples during rest are directly linked to the emergence of grid-cell-like neural code—a cellular mechanism representing the cognitive map (Constantinescu et al., 2016; Doeller et al., 2010).

Following rest, participants inferred relationships between compounds, each comprising three feature objects drawn from the learned feature dimensions. These compounds were positioned within a 2D task space defined by two feature dimensions. This 2D inference requires an internal model of the task space—i.e., a cognitive map.

We found that 2D inference performance was closely related to the feature knowledge acquired during the learning phase (*r* = 0.62, *P* = 9.567 × 10⁻⁵; **Fig. 4B**). To test whether this cognitive map is supported by a grid-like code, we analysed neural activity during the 2D inference task (**Fig. 4C**; see Methods). We focused on theta band (3–7 Hz) activity (Sebastijan Veselic et al., 2023), during the 3-second mental simulation period of each 2D inference trial (**Fig. 4A**), when participants mentally simulated an inferred trajectory in the 2D task space.

**Fig. 4.**
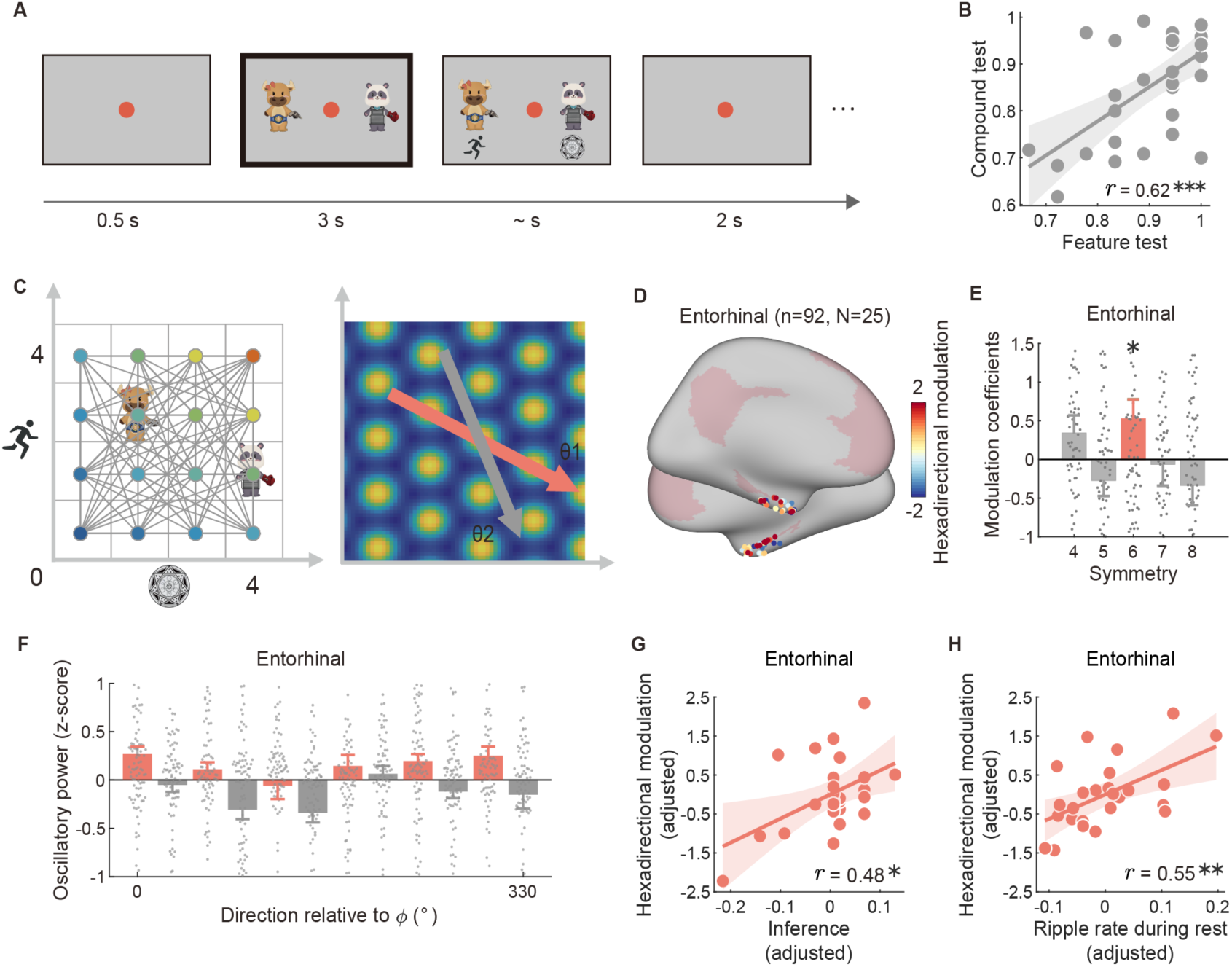
Grid-cell-like code in EC and mPFC theta oscillation during 2D inference. **A.** Core trial of 2D inference. Participants were tasked with comparing two compounds, either within one feature dimension or across two dimensions. Each trial began with a 0.5-second fixation period, followed by a 3-second presentation of the two compounds (highlight in black rectangle). Linking the two compounds mentally is akin to navigating through the mental space of all compounds, assuming that subjects have formed a 2D space. Subsequently, the decision cue is displayed below each compound; for example, comparing the ‘speed’ feature of the left compound to the ‘magic’ feature of the right one. Subjects were instructed to make their decision as quickly as possible.

In the entorhinal cortex (EC), we observed significant six-fold (hexadirectional) modulation in theta band activity (*t*_(89)_ = 2.10, *P* = 0.039), indicating the presence of grid-cell-like coding (**Fig. 4D–F**). We also examined the presence of grid-like codes within the DMN. Intriguingly, significant hexadirectional modulation was observed exclusively in the mPFC (*t*_(172)_ = 3.40, *P* = 8.547 × 10⁻⁴), with no significant results in other DMN regions (**Supplementary Fig. 3**, see also **Supplementary Fig. 4**). Control analyses confirmed that there was no selective six-fold modulation in other frequency bands—delta (1–3 Hz), alpha (7–12 Hz), beta (12–30 Hz), or gamma (30–60 Hz).

Importantly, the hexadirectional modulation in the EC was significantly correlated with inference performance, even after adjusting for memory performance (*r* = 0.48, *P* = 0.017; **Fig. 4G**). Moreover, this six-fold modulation was positively correlated with the hippocampal ripple rate during the preceding rest period, after controlling for the overall mean ripple rate across the entire task (*r* = 0.55, *P* = 0.005; **Fig. 4H**). These findings suggest that hippocampal ripples during rest are directly linked to the formation of grid-cell-like codes, supporting the formation of cognitive maps necessary for the inference tasks.

The trial ends with a 2-second inter-trial interval. **B.** The performance of the feature test before 2D inference is positively correlated with that of the compound test during 2D inference. **C.** Schematic representation of all inferred trajectories (the path connecting two compounds) on a 2D mental space (left panel). During the compound presentation period (black rectangle on panel A), subjects can mentally simulate the inferred trajectory on the 2D space, and if they do, we can look for the grid-cell-like code, based on the analysis of the hexadirectional modulation (right panel). The analysis of hexadirectional modulation is derived from observing grid-like firing patterns in grid cells (Doeller et al., 2010) and has been successfully detected in humans using fMRI, MEG and iEEG (Doeller et al., 2010; Kunz et al., 2015; Qu et al., 2024; Sebastijan Veselic et al., 2023; Staudigl et al., 2018). Each inferred trajectory has an angular direction (θ) and can be classified as aligned (red) or misaligned (grey) relative to the mean orientation of the grid-cell-like code (φ). Due to the hexagonal pattern of the grid cell, there would be a six-fold symmetry in the periodic change of neural activity if it is modulated by the grid code. **D.** Electrode contacts in the entorhinal cortex that exhibited hexadirectional modulation in theta band (3-7 Hz) activity. Each dot is a contact, colour coded based on the modulation strength. **E.** Selective six-fold modulation was observed in the entorhinal cortex, with no modulation found in other folds. Each dot represents one contact. **F.** The inferred trajectory (8) exhibits a six-fold modulation aligned to the mean orientation (φ) in the entorhinal cortex theta band activity. Effects are plotted separately for aligned (red) and misaligned (grey) trajectories. **G.** Positive correlation was observed between feature inference performance prior to the 2D inference phase and hexadirectional modulation in the entorhinal cortex measured during 2D inference, after controlling for memory performance. Each dot represents one subject. **H.** Hexadirectional modulation in the entorhinal cortex is also positively associated with the ripple rate during the preceding rest period, after controlling for the general mean ripple rate across the entire task duration. Error bars show SEM. * *P* < 0.05, ** *P* < 0.01, *** *P* < 0.001.

### Ripple-aligned mPFC activity during rest predicts later 2D inference performance

Building on the observed link between hippocampal ripples during rest and the emergence of grid-cell-like codes, we investigated whether cortical activity aligned with these ripples during rest is important for future inference abilities. Hippocampal ripples are among the most synchronous neuronal population patterns in the mammalian brain (Buzsáki, 2015), and interactions between the hippocampus and cortex during ripples are crucial for information transmission (Battaglia et al., 2011; Kaefer et al., 2022; Norman et al., 2021).

We focused on high-frequency broadband (HFB; 60–160 Hz, also known as high gamma) cortical activity, a robust marker of local neuronal spiking (Mukamel et al., 2005; Parvizi & Kastner, 2018; Ray et al., 2008). Specifically, we examined HFB activity across all cortical contacts during periods when the simultaneously recorded hippocampal ripples occurred (within ±250 ms of the ripple peak), referred to as *peri-ripple cortical activity during rest* (**Fig. 5B**), and assessed its impact on subsequent 2D inference performance.

**Fig. 5.**
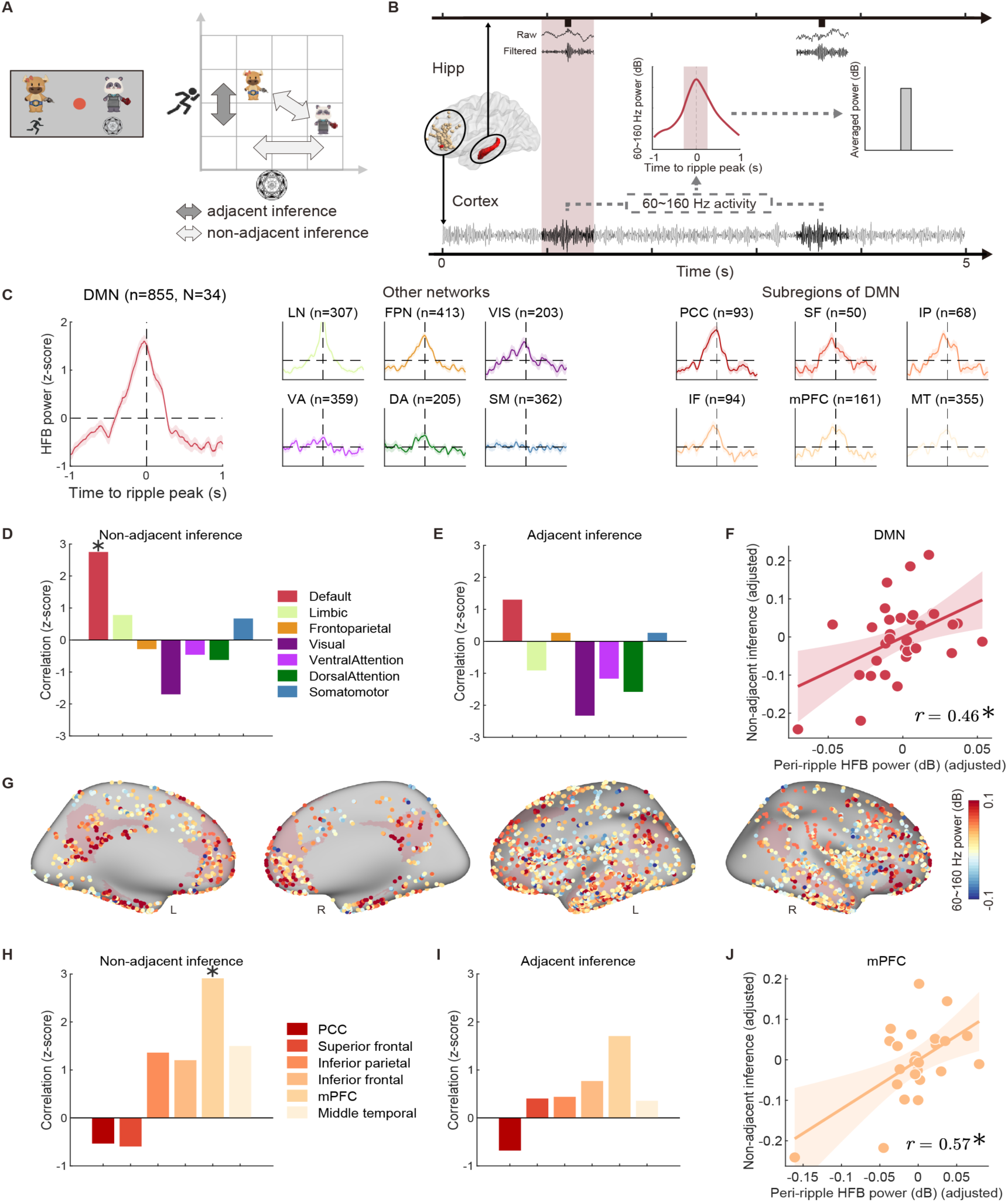
Hippocampal ripple-aligned high gamma activity in the mPFC during rest predicts subsequent 2D inference ability. **A.** During 2D inference, there are two types of trials based on whether they can be solved solely based on memory, while all probes concern compounds made of feature objects. In trials where the task is to compare the rank between two adjacent compounds within a feature dimension, this can be achieved by retrieving memories about the relative rank of the learned feature pairs; this is termed ‘adjacent inference’. Conversely, all other trials are termed ‘non-adjacent inference’, which involves inferring unobserved relationships between compounds. **B.** Analysis of peri-ripple high-frequency broadband (HFB, 60–160 Hz) activity. HFB power was specifically extracted around the ripple peaks (from -1s to +1s), with baseline correction applied by subtracting the mean power between -1s to -0.5s from the ripple peaks. Inset plots depicting the extracted peri-ripple HFB power (±250ms around the ripple peak). The zero time point was aligned with the ripple peaks. The high-frequency band activity shown is from Subject 36, channel BP05, recorded between 1365.85 s and 1370.85 s after the experiment start, which also shows the corresponding ripples during this period. **C.** Temporal profiles of the peri-ripple HFB (z-scored for visualization) within seven resting state brain networks and subregions of the DMN during the rest period. The letter ‘n’ indicates the number of contacts, and ‘N’ the number of subjects. **D.** Correlations between the averaged peri-ripple HFB power around the ripple peak (±250ms) in the DMN and other brain networks in relation to ‘non-adjacent inference’ performance. This analysis employed robust partial correlation, controlling for differences in premise knowledge (feature test accuracy). Correlation coefficients were converted to z-scores to visualize the differences across resting networks. **E.** Similar to panel D, but it is partial correlation in relation to ‘adjacent inference’. No significant correlation was found. **F.** Peri-ripple HFB power of the DMN during rest, predicted later ‘non-adjacent inference’ performance, after controlling for differences in premise knowledge. See also **Supplementary Fig. 5, 6**. The scatter plot represents partial regression. Each dot represents a single subject. **G.** Distribution of contacts within the DMN, with each contact colour-coded according to its peri-ripple HFB power strength. **H.** The partial correlation coefficients between the peri-ripple HFB power in mPFC, as well as other subregions within DMN, in relation to the ‘non-adjacent inference’ performance. Only mPFC was significant. **I.** Similar to panel H, but it is partial correlation in relation to ‘adjacent inference’ performance, no significant relationship was found. **J.** Peri-ripple HFB power in the mPFC during rest, predicted later ‘non-adjacent inference’ performance, after controlling for differences in premise knowledge. The scatter plot represents partial regression. Each dot represents a single subject. * *P* < 0.05. Abbreviation: HFB - high-frequency broadband; DMN - default mode network; LN - limbic network; VIS - visual network; FPN - frontoparietal network; VA - ventral attention network; DA - dorsal attention network; SM - somatomotor network; mPFC - medial prefrontal cortex; PCC - posterior cingulate cortex.

To distinguish inference from simple memory retrieval, we categorised the 2D inference trials into *adjacent inference* and *non-adjacent inference* (**Fig. 5A**). Adjacent inference trials involved comparing compounds that differed by only one rank in a feature dimension and could be solved by retrieving directly learned rank information. In contrast, non-adjacent inference trials involved comparing compounds with larger rank differences, requiring the integration of learned information to infer unobserved relationships.

We found that peri-ripple HFB activity in the DMN was significantly elevated compared to baseline periods (–1,000 ms to –500 ms relative to ripple peaks; see **Supplementary Fig. 5A**), indicating coordination between the DMN and hippocampal ripples during rest (**Fig. 5C**). Importantly, this peri-ripple HFB activity in the DMN exclusively predicted performance on non-adjacent inference trials (*r* = 0.46, *P* = 0.04; Bonferroni-corrected for all seven resting-state networks; **Fig. 5D** and **5F**), even after controlling for differences in prior knowledge. No significant relationship was found for adjacent inference performance (all *P* > 0.2; **Fig. 5E**). Notably, the hippocampal ripple rate during rest did not correlate with performance on either adjacent or non-adjacent inference trials (all *P* > 0.6; see **Supplementary Fig. 6B–C**), indicating that it is the ripple-aligned cortical activity in the DMN, rather than ripple occurrence alone, that predicts inference performance.

To ensure that these effects were specific to rest, we examined ripple-associated activity during the on-task inference period (from compound onset to response). We found that ripple rates did not differ between correct and error trials for either adjacent or non-adjacent inference (all *P* > 0.25; **Supplementary Fig. 6D–E**). Moreover, peri-ripple HFB activity in the DMN during the on-task period did not predict non-adjacent inference performance (*P* = 0.603; **Supplementary Fig. 6F**). These findings suggest that the predictive relationship between peri-ripple DMN activity and inference performance is specific to rest periods.

The DMN is thought to encode internal models of the world—cognitive map (Baldassano et al., 2018; Constantinescu et al., 2016; S. A. Park et al., 2021). To identify specific regions within the DMN involved in hippocampal–DMN interactions, we examined the relationship between peri-ripple HFB activity in each DMN subregion (**Fig. 5G**; see also **Supplementary Fig. 5B**) and non-adjacent inference performance. Intriguingly, only peri-ripple HFB activity in the mPFC was significantly associated with non-adjacent inference performance (*r* = 0.57, *P* = 0.022; Bonferroni-corrected for all six DMN subregions; **Fig. 5H** and **5J**), even after controlling for differences in prior knowledge. No such association was found for adjacent inference performance (all *P* > 0.09; **Fig. 5I**). This positive relationship persisted even after controlling for general HFB activity and hippocampal ripple rate during rest (see **Supplementary Fig. 5C–D**). Furthermore, this finding was validated using a more fine-grained brain atlas (Schaefer et al., 2018) and by analysing the entire brain, confirming that only peri-ripple mPFC activity during rest predicted later non-adjacent inference performance (see **Supplementary Fig. 7**). Together, these findings reveal an intriguing hippocampal-DMN dialogue during rest, particularly in the mPFC, which is important for future model-based inference ability.

## Discussion

Our study reveals that during offline periods, human hippocampal ripples interact with DMN activation, particularly within the mPFC, to align new information with a 2D conceptual map, supporting inference performance that extends beyond simple memory retrieval. These results complement recent MEG work showing that replay helps build cognitive maps (Ou et al., 2025), and provide the first direct evidence that hippocampal ripples are a key neuronal mechanism for aligning novel experiences with grid-like schema representations in the human brain.

During learning, we observed that ripple rates in the hippocampus were higher during ITI than feedback periods and increased progressively with accumulated experience. This increase in ripple activity positively predicted future feature-inference performance, implying that these bursts of activity help align discrete experiences as learning progresses. Similar observations in rodents show that ripples during brief pauses contribute to learning in physical space (Girardeau et al., 2009; Joo & Frank, 2018), grow with additional experience (Cheng & Frank, 2008; Igata et al., 2021; Pfeiffer, 2022), and facilitate spatial navigation (Pfeiffer, 2022; Roux et al., 2017). By using intracranial recordings, we extend these ripple-related processes from rodent spatial tasks to abstract conceptual learning in humans.

Beyond integrating individual feature objects, we found that hippocampal ripples during post-learning rest promote the formation of a compact 2D map in the EC and mPFC, characterised by a grid-like code. Grid representations have long been linked to stable encoding of physical spaces (Fyhn et al., 2007; Rowland et al., 2016; Wills et al., 2010), whereas abstract conceptual spaces may require more sophisticated reasoning (Constantinescu et al., 2016; Gattis & Holyoak, 1996; Schnall & Gattis, 2022). Previous work using fMRI or MEG has identified grid-like patterns in non-spatial cognitive maps (Bao et al., 2019; Constantinescu et al., 2016; Liang et al., 2024; Ou et al., 2025; S. A. Park et al., 2021). Our data go further, providing direct evidence that hippocampal ripples align new experiences with a grid code in conceptual space.

Our iEEG approach captured rapid temporal dynamics and simultaneous hippocampal– cortical activity, identifying ripples as a crucial substrate for grid codes in the EC. In rodents, grid cells receive inputs from hippocampal place cells, which replay spontaneously during ripples (Foster & Wilson, 2006; Pavlides & Winson, 1989; Wilson & McNaughton, 1994). This replay may similarly align new experiences with human EC grid representations. While MEG data indicate that replay underlies grid-code emergence in resting humans (Ou et al., 2025), our findings provide the first direct demonstration that hippocampal ripples align conceptual information with a grid schema.

Interestingly, while ripples during rest predicted the later appearance of grid codes, ripples alone did not directly correlate with inference accuracy. This suggests that robust inference demands further refinement or integration of the conceptual map. Because ripples are highly synchronous events, bidirectional interactions between the hippocampus and other brain regions are likely crucial for more sophisticated reasoning (Norman et al., 2019, 2021; Pedrosa et al., 2022; Verzhbinsky et al., 2024). Here, aligning neural data from 3,136 cortical contacts (902 in the DMN) to hippocampal ripples across 38 subjects enabled us to characterise transient, brain-wide activity tied to ripple events. Although six of the seven resting-state networks (excluding the somatomotor network) showed heightened activity during ripples, only DMN activity in the mPFC during rest significantly predicted subsequent inference performance. This was especially true for non-adjacent inference trials that required an internal map of the task space rather than simple memory retrieval.

The DMN is believed to encode an internal model of the world (Constantinescu et al., 2016; Menon, 2023; S. A. Park et al., 2021; Yeshurun et al., 2021), with the mPFC integrating significant inputs from the hippocampus (Chao et al., 2020; Eichenbaum, 2017; A. J. Park et al., 2021; Peyrache et al., 2009). Ripple-coupled activation within the DMN has been observed during episodic memory recollection (Norman et al., 2021), and our study highlights its role in aligning conceptual representations for inference. In rodents, the mPFC quickly consolidates new knowledge with existing frameworks (Farzanfar et al., 2023; Tse et al., 2007) and forms memory engrams through hippocampal input (Kitamura et al., 2017). The mPFC also exerts top-down control over the hippocampus to adapt to different contexts (Eichenbaum, 2017; Malik et al., 2022; Miller & Cohen, 2001). Thus, while ripples anchor new experiences to a grid code, the mPFC may be required to fully integrate and apply this map for complex inference.

In summary, our results show that hippocampal ripples, in coordination with the DMN, align new experiences with grid-like conceptual schemas, transforming discrete learning events into structured knowledge that supports flexible inference. By bridging animal findings on ripples and replay with human neuronal recordings, we demonstrate how non-spatial conceptual tasks draw upon grid representations for reasoning. This highlights the critical role of hippocampal–cortical coordination through ripple events in human cognition.

## Methods

### Participants

A total of 42 patients were initially included in this study; four did not meet the learning criteria, resulting in 38 patients (12 females; demographics and electrode contacts coverage information are in **Tables S1 and S2**) for formal data analysis. All subjects were with medicine-resistant epilepsy underwent intracranial depth electrode implantation as part of their pre-surgical evaluation at Beijing Sanbo Brain Hospital of Capital Medical University. The electrode implantation locations were determined solely based on clinical evaluation. All experimental procedures were approved by the Ethics Committee of the Sanbo Brain Hospital of Capital Medical University, SBNK-YJ-2023-002-01.

### Task

#### Overview

The experiment was conducted in a clinical setting using PsychoPy (Peirce et al., 2019) (https://psychopy.org/) software on a Windows 7 system. It consisted of several task sessions: *premise learning*, *feature test*, and *2D inference*. Each premise learning session lasted approximately 6 minutes, followed by a feature test. Participants who performed well on the *feature test* proceeded to the subsequent task. Following this, participants had a 5-minute resting period with their eyes open. After the rest, they underwent feature test again. Finally, the 2D inference task, lasting approximately 17 minutes, concluded the experiment. On average, the entire experiment completed within 1 hour.

#### Premise learning

Participants were instructed to learn the hierarchical ranks within three feature dimensions: ‘speed’, ‘power’, and ‘magic’, each comprising of four objects, as illustrated in **Fig. 1A**. These objects were assigned attributes of ‘ability rank’ from 1 to 4 within each dimension, irrespective of real-world associations. As depicted in **Fig. 2A**, following a 0.5-second fixation period, two objects were simultaneously presented, accompanied by dimension cues indicating their respective dimensions, prompting participants to determine ‘which one is higher?’. The objects pairs remained on screen for no longer than 3 seconds until a response was made. Upon selection, the chosen object was indicated by a white rectangle, and the response outcome (correct or error) was displayed for 3 seconds. Participants learned the rank of these objects through feedback. A 2-second inter-trial interval followed before the next trial began. During the premise learning session, participants encountered a total of 9 adjacent pairs (3 for each dimension, totalling 3 dimensions), which were intermixed and repeated 5 times, leading to 45 trials in total. Participants were assessed on their performance in the *feature test*, achieving an average accuracy score of 0.88 ± 0.12 (mean ± SD) before the rest period. If participants performed poorly in the *feature test*, additional *premise learning* sessions were conducted. Each session involved a reduced number of trials, with 9 trials in total (1 for each pair). This process was repeated until participants successfully passed the feature test.

#### Feature test

During the feature test session, participants were assessed on their memory performance regarding pairwise rank relationships and their capability to infer relationships spanning more than one rank within a single feature dimension. This involved the simultaneous presentation of two distinct objects within the same feature dimension after a 0.5-second fixation, with participants required to determine which one ranked higher, without any time limitation. To prevent a learning effect, no feedback was provided after their decision. Subsequently, a 2-second inter-trial interval followed. The feature test session comprised 9 trials for memory testing (involving adjacent pairs) and 9 trials for feature inference testing (involving non-adjacent pairs), all of which were intermixed. The same feature test session was also administered after the rest period as a post-rest feature test.

### 2D inference

Participants who demonstrated proficiency in ranking features were directed to a compound test, wherein they performed inference based on compounds composed of these feature objects. Each compound consisted of three feature objects from three distinct feature dimensions. While all three feature dimensions were learned, only two of them were task-relevant during the two-dimensional (2D) inference task. The remaining dimension was randomly chosen for each subject to control visual variability. As depicted in **Fig. 4A**, following a brief 0.5-second fixation, two compounds were simultaneously presented for 3 seconds. This period, termed the mental simulation period, allowed participants to mentally trace the trajectory between the two compounds in a 2D task space. After this period, decision cues appeared below each compound, directing participants to compare specific features. For instance, they might be prompted to compare the ‘speed’ feature of the left compound to the ‘magic’ feature of the right one. Participants were instructed to make their decision as swiftly as possible. Each trial concluded with a 2-second inter-trial interval. Compound pairs were selected to cover more angles and ensure equal coverage for each location as possible. There were 120 trials in total during the 2D inference session. In trials where the task is to compare the rank between two adjacent compounds within a feature dimension, this can be achieved by retrieving memories about the relative rank of the learned feature pairs; this is termed ‘adjacent inference’, and resulted in 40 trials. Conversely, all other trials are termed ‘non-adjacent inference’, which involves inferring unobserved relationships between compounds.

### Intracranial EEG recordings

Electrophysiological recordings were conducted utilising a Blackrock clinical monitoring system with a sampling rate of 2000 Hz and a bandpass filter ranging from 0.3 Hz to 500 Hz. Depth electrodes featured a diameter of 0.8 mm and a height of 2 mm, with a 3.5 mm separation between the centres of adjacent electrode contacts.

### Electrode anatomical localisation

The MRI data initially underwent segmentation using FreeSurfer (Fischl, 2012) to distinguish between white and grey matter. Subsequently, the postimplant CT scan was aligned with the preimplant MRI anatomical scan using Brainstorm 2023 (Tadel et al., 2011). Manual labelling of electrode locations was performed within Brainstorm. For group-level analyses, SUMA was utilised to resample and standardize cortical templates (Saad & Reynolds, 2012), while iELVis (Groppe et al., 2017) facilitated visualization of all electrode contacts on a single cortical template, either ‘‘fsaverage’’ or ‘‘MNI305’’. The assignment of atlas labels to electrode contacts was conducted in the native space of each brain. Electrode contacts located more than 3 mm from the cortical ribbon were excluded from further analysis. Furthermore, each electrode contact was assigned to a functional-connectivity-based atlas (Yeo et al., 2011). Locations within the entorhinal cortex were delineated using the Julich-Brain atlas (Amunts et al., 2020). Cortical recording contacts, organised as bipolar pairs, were categorized based on their atlas label, referencing the first contact of each bipolar pair.

### Preprocessing and analysis of sEEG

Data analysis was conducted in MATLAB 2022b (MathWorks Inc., Natick, MA) utilising EEGLAB v2022.1 (Delorme & Makeig, 2004), alongside custom-developed analysis code. The raw sEEG signal, sampled at 2000 Hz, underwent down sampling to 1000 Hz, followed by statistical inspection to identify noisy or corrupted channels for exclusion from further analyses. Channels exhibiting voltage values, voltage derivative, or RMS in the 99th percentile exceeding 5 SD compared to other contacts were flagged for exclusion after visual assessment. Next, the signal underwent notch filtering to eliminate 50 Hz power line interference and its harmonics (100 Hz, 150 Hz, etc.) using a zero-lag linear-phase Hamming-windowed FIR band-stop filter (3 Hz wide). The filtered healthy iEEG signals were then converted into bipolar derivations, pairing adjacent recording contacts on the same electrode probe. Recording contacts within the hippocampus were paired with nearby electrode contacts in white matter.

### Hippocampal ripples detection

The detection of hippocampal ripples was achieved through established procedures (A. A. Liu et al., 2022; Norman et al., 2019; Stark et al., 2014; Vaz et al., 2019). Electrode contacts positioned within 3 mm of the hippocampus were selected, and a nearby white-matter contact’s reference signal was subtracted to reduce common noise. The intracranial EEG (iEEG) signals were bandpass filtered between 70-180Hz (zero-lag linear-phase Hamming windowed FIR filter) and subjected to a Hilbert transform to compute instantaneous analytic amplitude. Ripple events were identified based on their amplitude exceeding 4 standard deviations (SD) above the mean. The duration of these events extended until ripple power fell below 2D, with the ripple peak aligned to the trough closest to the peak. Candidate ripple events were constrained to durations between 20 ms to 200 ms, with a 30 ms interval between events. Artifacts were minimized through control detection on the common average signal, typically affecting around 3 ± 2 % of the ripples detected in each contact. Pathological events, such as interictal epileptic discharges (IEDs) and pathological high-frequency oscillations (HFOs), were excluded. Finally, the identified ripples underwent visual inspection in both time and frequency domains after averaging within contacts, with un-normal contacts excluded from further analysis.

### High-frequency broadband signal

In the current study, the high-frequency broadband (HFB) signal, also referred to as high-gamma, is defined as the mean normalized power within the frequency range of 60–160 Hz. Previous research has established that activity within this frequency band serves as a robust electrophysiological marker of local neuronal spiking (Mukamel et al., 2005; Parvizi & Kastner, 2018; Ray et al., 2008). To compute HFB power, zero-lag linear-phase Hamming-windowed FIR filters of order 660 were applied to segment the signal into 10 Hz bands within the 60-160 Hz range. Subsequently, the normalized and 1/f corrected analytic amplitude was calculated using a Hilbert transform. The resulting amplitude was then converted to decibels (dB) using the formula 10*log_10_(Amplitude) for statistical analysis.

### Mixed-effects analysis

Mixed-effects analyses were carried out using the LME4 package (Bates et al., 2018) in R (Team, 2013). Data were fitted with a random intercept model, incorporating relevant fixed factors and a random factor of ‘Contact’ nested within ‘Participant’ to account for varying contributions across participants. In examining the influence of response outcomes on ripple rate during learning (referencing **Fig. 2C**), the model was structured as follows:

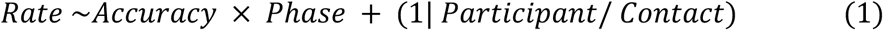

Here, accuracy (correct/error) and phase (feedback/interval) are categorical variables with two levels each, while ripple rate is continuous. For analysing the interaction between learning process and response accuracy on ripple rates (**Fig. 2D**), the models were formulated as follows:

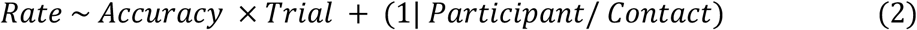

Where trial number is a continuous variable. In investigating the effect of ripples or peri-ripple HFB on performance during 2D inference sessions, the models were formulated as follows:

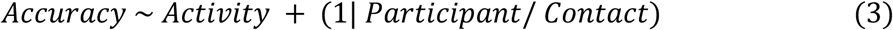

Here, accuracy is represented by binary variables (0 and 1). Main effects were tested using Type III ANOVA test, with degrees of freedom computed using the Kenward-Roger (KR) method. Post hoc comparisons were conducted using the emmeans package (RV Lenth, 2020), and corrections for multiple comparisons were applied using the Holm method (Holm, 1979).

### Hexadirectional modulation analysis

We conducted hexadirectional modulation analyses following established methods (Constantinescu et al., 2016; Doeller et al., 2010; Maidenbaum et al., 2018; S. A. Park et al., 2021). This approach enabled the detection of grid cell signals across various imaging modalities, including fMRI (Constantinescu et al., 2016; Doeller et al., 2010; Kunz et al., 2015), iEEG (Maidenbaum et al., 2018; Sebastijan Veselic et al., 2023), and MEG (Staudigl et al., 2018), extending beyond the microscopic level. In essence, all compounds under scrutiny were mapped onto a 2D plane based on their attributes. Trajectories’ angles (θ) between pairs of compounds within individual trials were computed, determining whether they aligned or misaligned with the orientation (Φ) characterizing grid cells, thereby influencing neural activity.

To execute this, half of the data (e.g., trials 1, 3, 5, etc.) were utilised to estimate the grid orientation within theta band (3 – 7 Hz, zero-lag linear-phase Hamming windowed FIR filter, 0.5 Hz transition band) activity, which was then validated using the remaining trials. Grid orientation estimation involved modelling neural activity in the theta band using a general linear model (GLM1) with cosine and sine regressors:

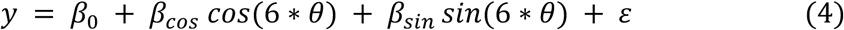

The resulting weights from these regressors, *β_cos_* and *β_sin_*, were utilised to calculate the grid orientation (*Φ*) using the formula:

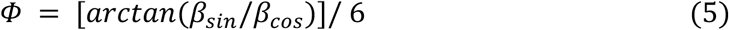

Subsequently, a second general linear model (GLM2) was employed to assess whether theta band activity increased as participants moved more aligned with the grid orientation *Φ*. This model quantified the extent of sixfold rotationally symmetric modulation of theta activity by movement direction:

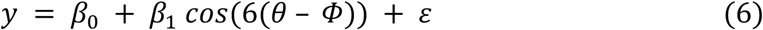

Higher *β_1_* values indicated greater neural activity when subjects moved closer to the preferred orientation *Φ* (and conversely, lower power when moving closer to nonpreferred orientations in between). The factor 6 accounted for the sixfold rotational symmetry, as depicted in Figure 4C. Additionally, other symmetries (such as 4, 5, 7, and 8) were also tested for contrast.

To mitigate trial selection bias, the process was repeated with the training and testing data sets reversed, and the results from both procedures were averaged. For an intuitive perspective on the sixfold modulation effect, theta band activity was then categorized into alignment and misalignment based on whether the trajectory (*θ*) aligned with the estimated grid orientation (*Φ*).

## Acknowledgment

Conceptualization, Y.L., Z.X., J.O., Y.Q., and T.B.; Investigation, Z.X., X.W., J.Z., L.H., X.H., Y.L., and T.B.; Writing – Original Draft, Z.X., Y.L.; Writing – Review & Editing, Z.X., Y.L., L.H., and T.B. This study is supported by the National Science and Technology Innovation 2030 Major Programme (2022ZD0205500), the National Natural Science Foundation of China (32271093), the Beijing Natural Science Foundation (Z230010, L222033), and the Fundamental Research Funds for the Central Universities.

## Data Availability

The processed data, including detected ripple events, theta band activity, high-frequency broadband activity, and corresponding behavioural data, will be made available at https://doi.org/10.5281/zenodo.12794280 upon publication.

## Code Availability

The analysis code will be publicly accessible at https://gitlab.com/liu_lab/ripple-representation-learning upon publication.

## Supplementary Information

**Supplementary Fig. 1.**
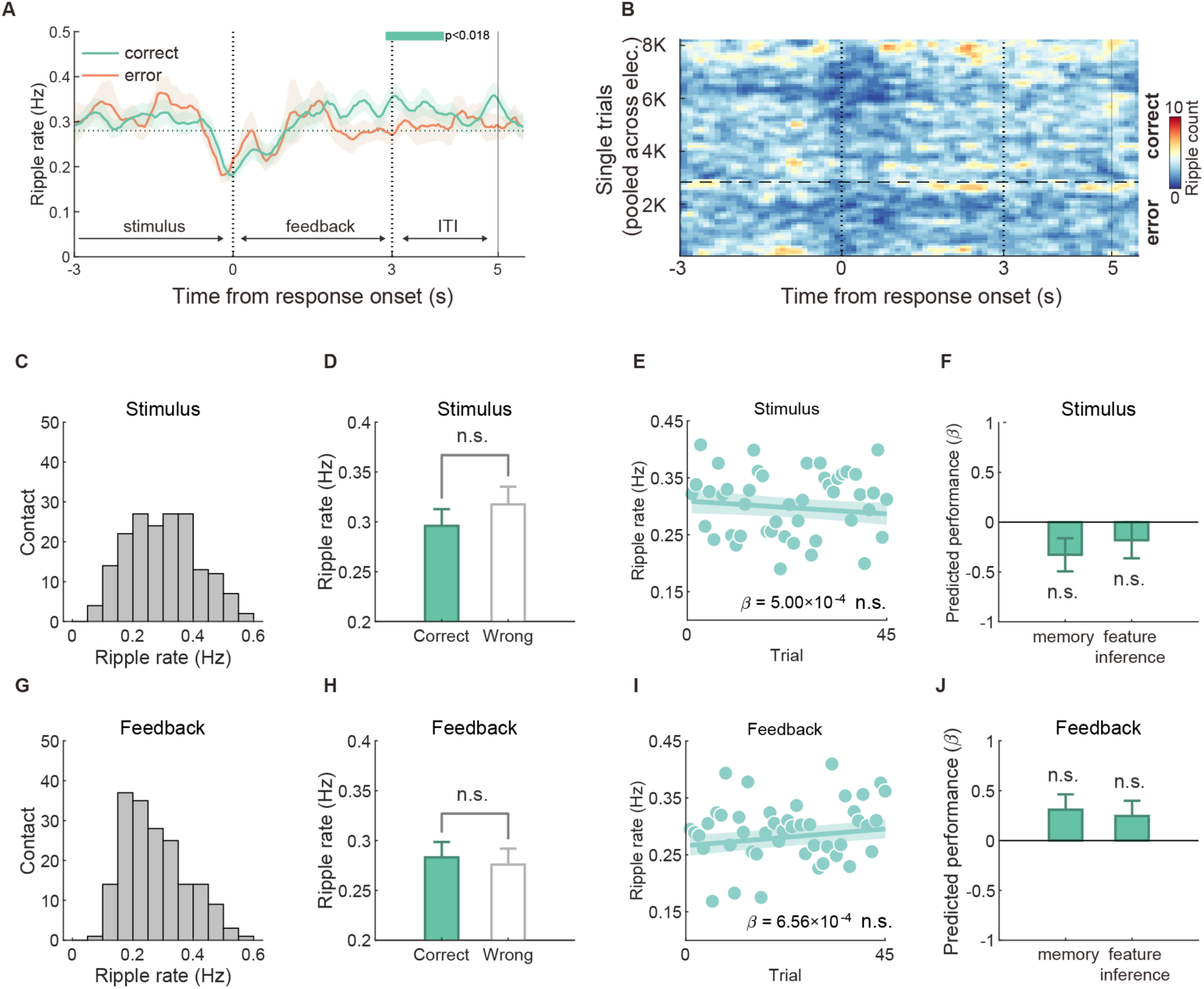
Hippocampal ripples during stimulus or feedback periods did not increase with learning, nor were they related to future behaviour. **A.** Group-level averaged peri-event time histograms (PETH) of ripples, time-locked to the onset of response, categorized into trials with correct responses and error responses. The green bar indicates significant time bins between correct and error trials, corrected for multiple comparisons using a cluster-based permutation test. **B.** Density plot depicting hippocampal ripple probability for correct and error response trials, pooled across all electrodes. Ripple density was computed in bins of 100 ms × 100 trials and smoothed using a 3-bins-wide Gaussian filter for visualization. **C.** Distribution of ripple rates during stimulus presentation. **D.** Ripple rates during stimulus presentation did not show significant differences between correct and error response trials. **E.** Ripple rates during stimulus presentation in correct response trials did not exhibit significant changes over the course of learning (number of learning trials). **F.** The Learn-R (Learning-Increased Ripple) during stimulus presentation was not predictive of feature inference or memory performance. **G.** Distribution of ripple rates during feedback presentation. **H.** Ripple rates during feedback presentation did not significantly differ between correct and error response trials. **I.** Ripple rates during feedback presentation in correct responded trials did not show significant change over the course of learning. **J.** Similarly, the Learn-R (Learning-Increased Ripple) during the feedback presentation was not predictive for feature inference or memory performance. Each dot represents one subject. Error bars show SEM. n.s., not significant.

**Supplementary Fig. 2.**
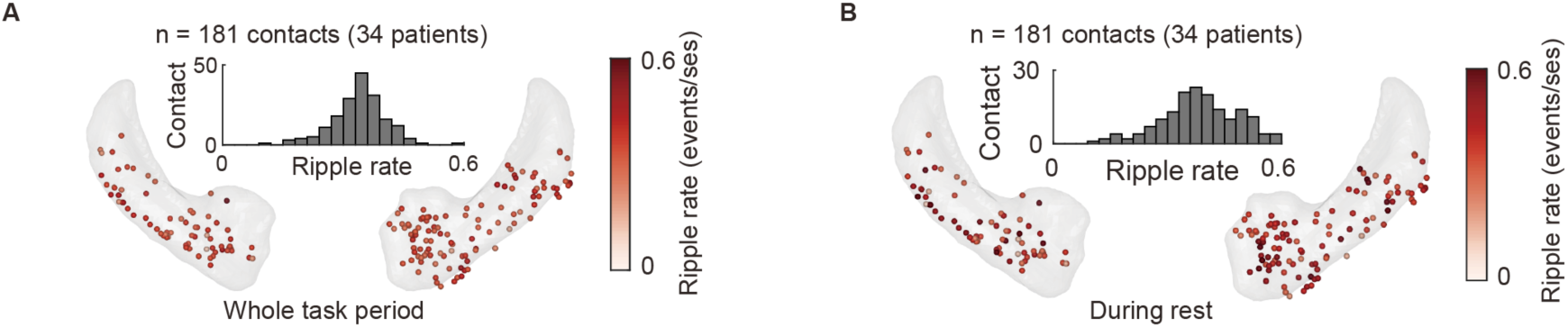
Distribution of hippocampal ripple rates during both rest periods and the entire task. **A.** Hippocampal ripple rates across all contacts throughout the task, with a histogram inset. **B.** Similar to panel A, but focusing on the 5-minute rest period.

**Supplementary Fig. 3.**
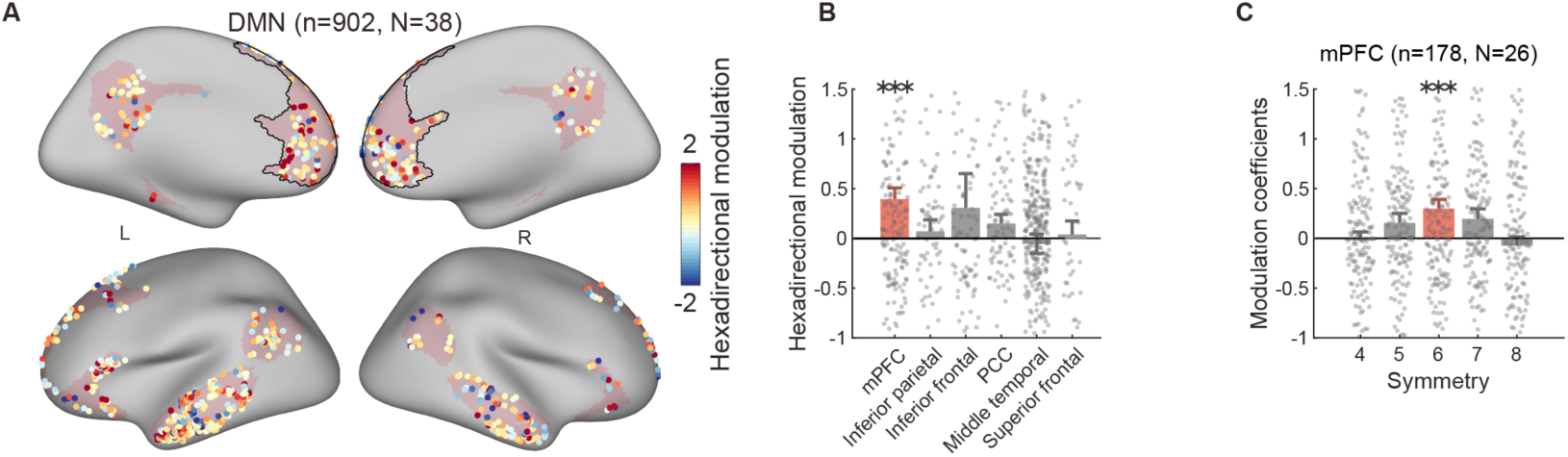
Grid-like code in DMN. **A.** Electrode contacts within the DMN are colour-coded to represent the strength of hexadirectional modulation in the theta band activity. The mPFC is highlighted with a black contour. **B.** Across all sub-regions within the DMN, hexadirectional modulation was exclusively observed in the mPFC. **C.** Selective six-fold modulation was observed in the mPFC, with no modulation found in other folds. Error bars show SEM. ***P < 0.001. Abbreviation: DMN - default mode network; mPFC - medial prefrontal cortex; PCC

**Supplementary Fig. 4.**
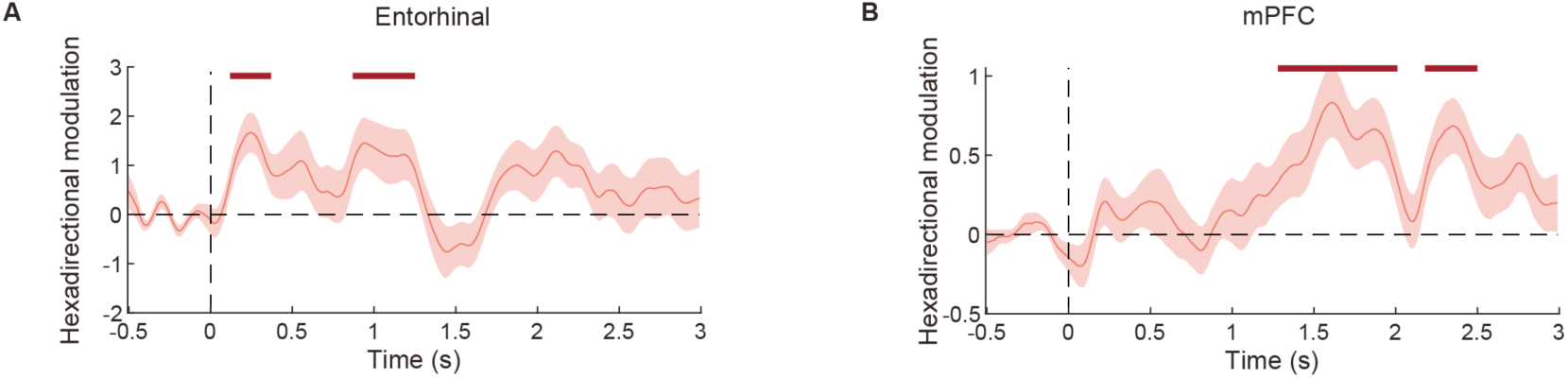
Temporal dynamics of hexadirectional modulation in the EC and mPFC. **A.** Analysis of the temporal characteristics of hexadirectional modulation in theta-band activity within the entorhinal cortex (EC) during mental simulation. Time zeros were locked to the onset time of two compounds. Hexadirectional modulation coefficients were computed for each time point in each contact. **B.** Similar to panel A, but focusing on the medial prefrontal cortex (mPFC). Error bars show SEM. The red flag bar highlights significant time bins, identified through cluster-based permutation tests (*P* < 0.05), compared to the baseline period (-0.5s to 0s).

**Supplementary Fig. 5.**
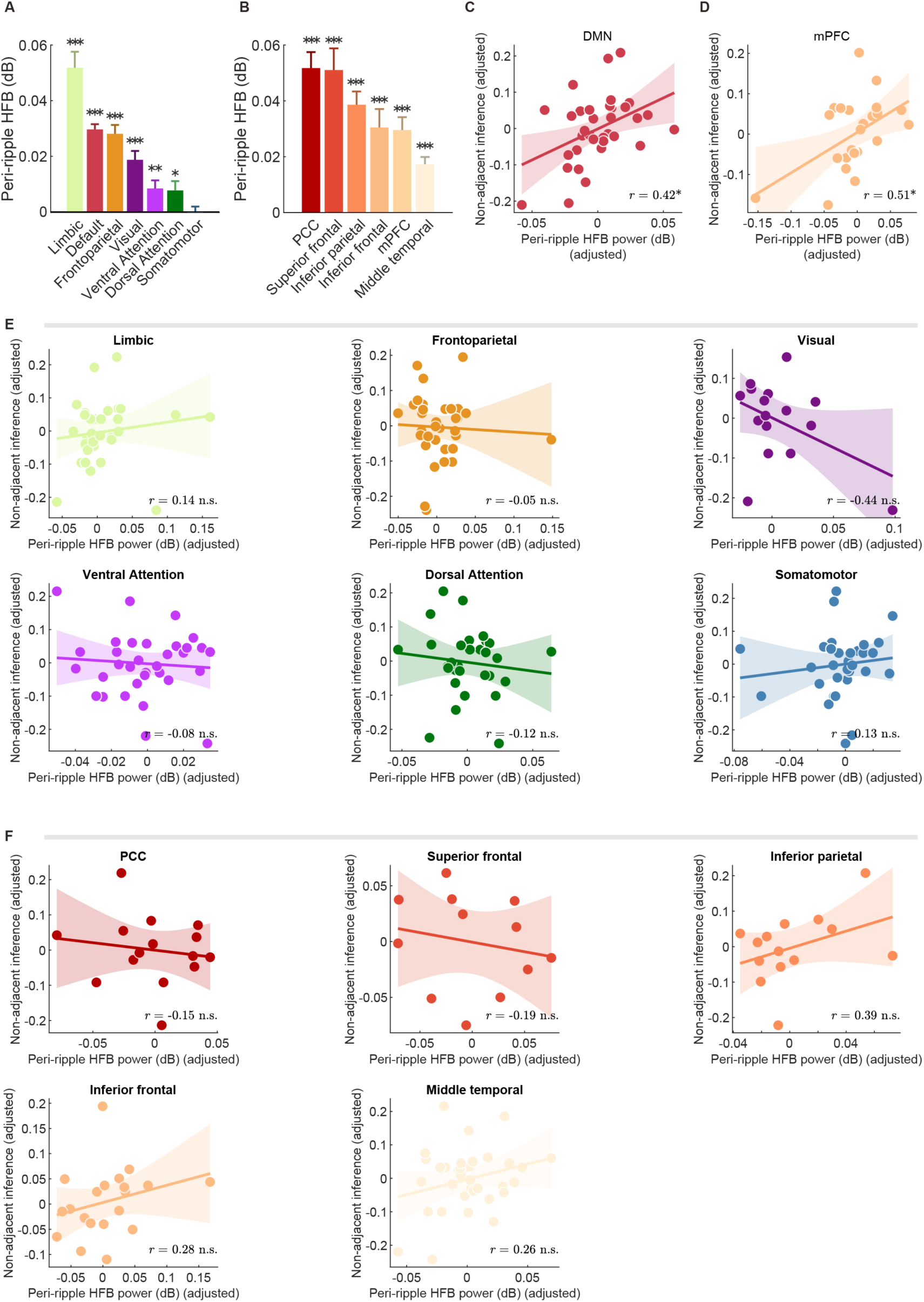
Control analysis of the relationship between peri-ripple HFB during rest and non-adjacent inference. **A.** The averaged peri-ripple HFB power around ripple peak (±250ms) for six out of seven rest-state brain networks were higher than the baseline period (-1000 ms to -500 ms relative to ripple peaks), except for the somatomotor network. **B.** Similar to panel A, but within sub-regions of the Default Mode Network (DMN). **C.** Correlations between the averaged peri-ripple HFB power around the ripple peak (±250ms) in the DMN and other brain networks in relation to ‘non-adjacent inference’ performance. This analysis employed robust partial correlation, controlling for differences in premise knowledge (feature test accuracy), general HFB activity, and ripple rate during rest. **D.** Peri-ripple HFB power of the DMN during rest, predicted later ‘non-adjacent inference’ performance, after controlling for differences in premise knowledge, ripple rate and general HFB during rest. Each dot represents a subject. The solid line indicates the linear fit. **E.** Scatter plot illustrating the correlations shown in Fig. 5D, depicting the relationship between non-adjacent inference performance and peri-ripple HFB activity within resting-state networks during rest. **F.** Similar to panel E, but focused on sub-regions of the DMN. Each dot indicates one subject. Error bars show SEM. * *P* < 0.05, ** *P* < 0.01, *** *P* < 0.001. Abbreviation: HFB - high-frequency broadband; mPFC - medial prefrontal cortex; PCC - posterior cingulate cortex.

**Supplementary Fig. 6.**
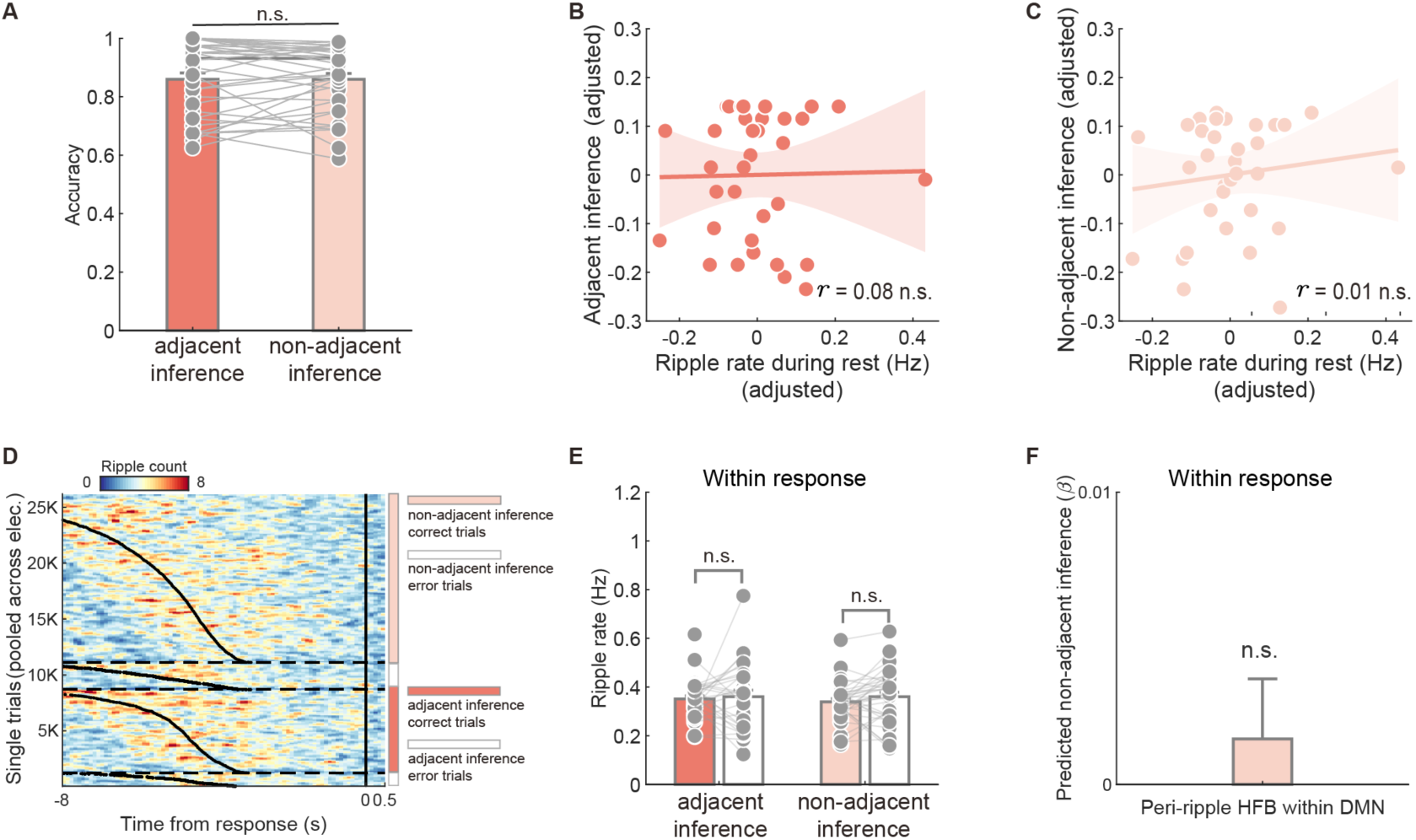
Compound-based inference is not related to hippocampal ripple activity during rest, nor to on-task hippocampus-DMN dialogue. **A.** The performance of ‘adjacent inference’ and performance of ‘non-adjacent inference’ in compound test were comparable, without significant difference. **B.** The performance of ‘adjacent inference’ was not correlated with ripple rate during rest, after controlling for the general mean ripple rate across the entire task duration. **C.** The performance of ‘non-adjacent inference’ was not correlated with ripple rate during rest, after controlling for the general mean ripple rate across the entire task duration. **D.** The hippocampal ripple density plot illustrates the variations in ripple probability before responses are made in ‘adjacent inference’ and ‘non-adjacent inference’ trials, distinguishing between trials with correct and incorrect responses. Trials were pooled together across all electrodes and sorted according to reaction time (black curve). Ripples’ density was computed in bins of 100 ms × 100 trials, smoothed using a 3-bins-wide Gaussian filter for visualization purposes. **E.** Ripple rates during the 2D inference period (from compounds onset to response) does not differ between correct and error trials in both ‘adjacent inference’ and ‘non-adjacent inference’ trials. **F.** High-frequency broadband (HFB) activity aligned with ripples in the DMN does not predict ‘non-adjacent inference’ during the 2D inference period. Error bars show SEM. n.s., not significant. Abbreviation: DMN - default model network.

**Supplementary Fig. 7.**
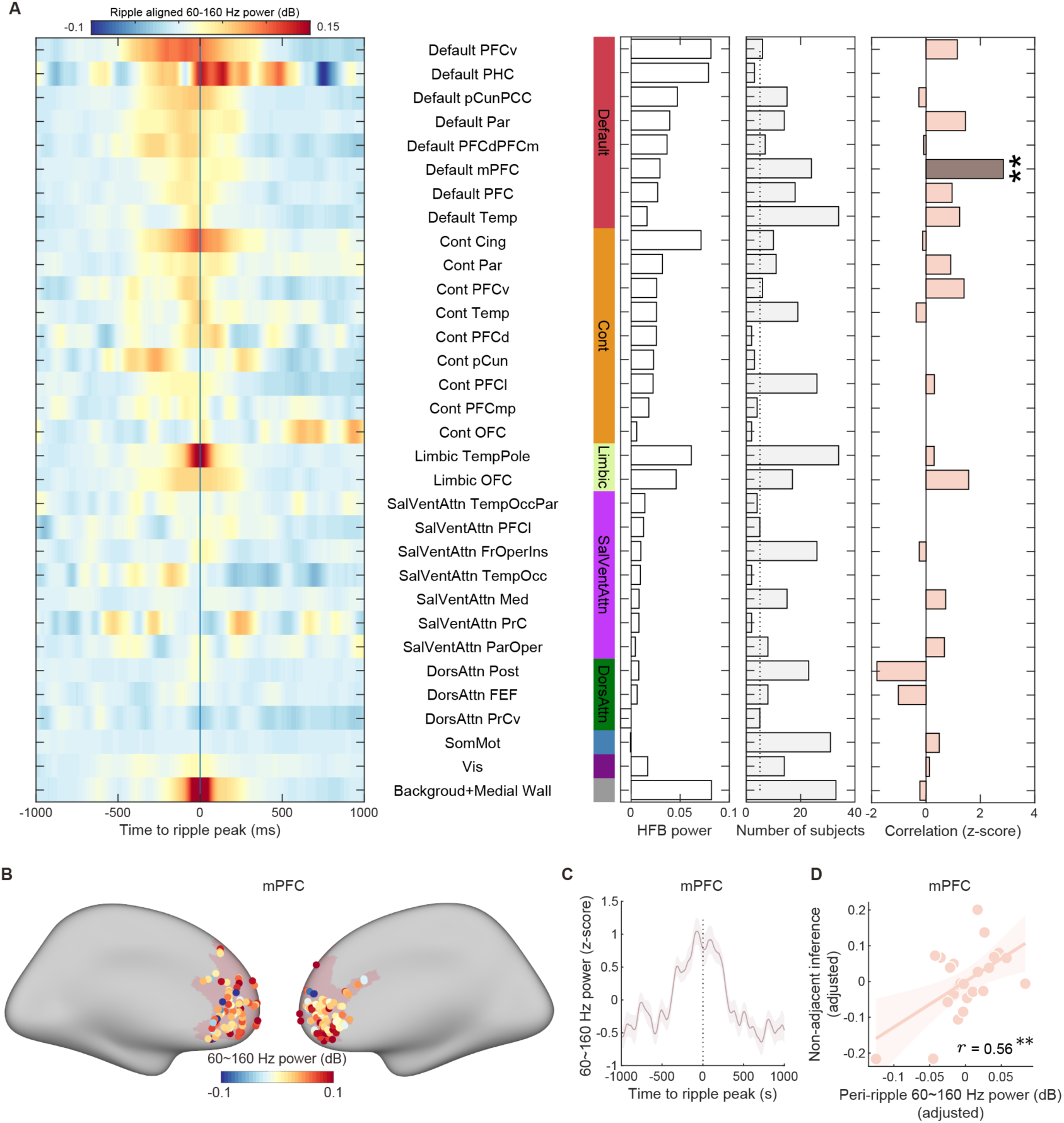
Brain-wide activity aligned with hippocampal ripples and its impact on 2D inference. **A**. Ripple-aligned power in the 60-160 Hz range spans the entire brain. Sub-regions were defined using the *S7* network atlas (Schaefer et al., 2018). The mPFC represents the medial part of the *PFC* within the default network. These regions were manually separated from the *PFC* and *PFCdPFCm* in the *S7* network atlas, with no overlap among sub-regions in this study. Bar plots display averaged peri-ripple HFB power (±250 ms), along with the number of subjects and correlation coefficients (z-scored) between peri-ripple HFB power and non-adjacent inference. The correlation analysis employed robust partial correlation, controlling for differences in premise knowledge (feature test accuracy). Dark colours in correlation bar plots indicate significant findings, with correlation coefficients shown only for sub-regions that included more than five subjects. **B.** Distribution of contacts within the mPFC, with each contact colour-coded according to its peri-ripple HFB power strength. **C.** Waveform of 60∼160 Hz power around ripple peaks in the mPFC. The power strength was normalized across time. **D.** Correlation between peri-ripple HFB in the mPFC and non-adjacent inference after controlling for differences in premise knowledge. The scatter plot represents partial regression. Each dot represents a single subject. ** *P* < 0.001.

**Table S1.** Demographic information for each patient, related to the Methods.

| Subject # | Gender | Age | RPM score | MMSE | Education years | Handedness |
| --- | --- | --- | --- | --- | --- | --- |
| SUB01 | Male | 15 | 58 | 30 | 9 | Right |
| SUB02 | Male | 36 | 32 | 27 | 12 | Right |
| SUB03 | Female | 30 | 49 | 27 | 16 | Right |
| SUB04 | Female | 19 | 47 | 30 | 9 | Right |
| SUB05 | Male | 15 | 49 | 30 | 12 | Right |
| SUB06 | Male | 29 | 32 | 30 | 16 | Right |
| SUB07 | Male | 22 | 41 | 30 | 16 | Right |
| SUB08 | Male | 35 | 41 | 30 | 6 | Right |
| SUB09 | Male | 25 | 34 | 29 | 12 | Right |
| SUB10 | Male | 22 | 52 | 30 | 16 | Right |
| SUB11 | Female | 23 | 55 | 30 | 16 | Right |
| SUB12 | Female | 14 | - | 29 | 9 | Right |
| SUB13 | Male | 19 | 47 | - | 16 | Right |
| SUB14 | Male | 15 | 14 | 29 | 12 | Right |
| SUB15 | Male | 24 | 47 | 30 | 16 | Left |
| SUB16 | Female | 33 | 50 | 29 | 16 | Right |
| SUB17 | Male | 49 | 53 | 30 | 16 | Right |
| SUB18 | Male | 18 | - | 29 | 12 | Right |
| SUB19 | Male | 30 | 46 | 30 | 12 | Right |
| SUB20 | Male | 19 | 56 | 30 | - | Right |
| SUB21 | Male | 30 | 52 | 30 | 12 | Right |
| SUB22 | Male | 30 | 57 | 30 | 16 | Right |
| SUB23 | Female | 14 | 52 | 30 | 9 | Right |
| SUB24 | Male | 26 | 22 | 30 | 12 | Right |
| SUB25 | Female | 42 | 33 | 28 | 9 | Right |
| SUB26 | Male | 24 | 49 | 26 | 16 | Right |
| SUB27 | Female | 26 | 50 | 30 | 16 | Left |
| SUB28 | Male | 18 | 52 | 30 | 12 | Right |
| SUB29 | Male | 15 | 54 | 30 | 9 | Right |
| SUB30 | Male | 16 | - | 30 | 12 | Right |
| SUB31 | Male | 26 | 55 | 30 | 16 | Right |
| SUB32 | Male | 27 | 56 | 30 | 16 | Right |
| SUB33 | Female | 54 | 44 | 30 | 16 | Right |
| SUB34 | Male | 19 | 45 | 28 | 9 | Right |
| SUB35 | Male | 34 | 34 | 29 | 16 | Right |
| SUB36 | Female | 16 | 56 | 30 | 12 | Right |
| SUB37 | Female | 13 | 58 | 30 | 9 | Right |
| SUB38 | Female | 37 | 31 | 30 | 16 | Right |
| M±SD | 12/26 | 25±9.8 | 46±10.7 | 29±1.0 | 13±3.1 | 2/36 |

Note: The RPM score represents the raw score of Raven's Progressive Matrices, and the MMSE score represents the raw score of the Mini-Mental State Examination. The '-' indicates that data for this item are missing.

**Table S2.** Distribution of recording contacts for each patient, related to the Methods.

| Subject # | Total analysed | Hippocampal contacts | Within Entorhinal | Within DMN | Within mPFC |
| --- | --- | --- | --- | --- | --- |
| SUB01 | 85 | 4 | 3 | 28 | 12 |
| SUB02 | 83 | 3 | 3 | 29 | 2 |
| SUB03 | 90 | 6 | 0 | 18 | 5 |
| SUB04 | 30 | 0 | 0 | 3 | 0 |
| SUB05 | 80 | 11 | 11 | 20 | 0 |
| SUB06 | 60 | 8 | 4 | 22 | 0 |
| SUB07 | 97 | 2 | 3 | 16 | 7 |
| SUB08 | 57 | 0 | 0 | 23 | 13 |
| SUB09 | 71 | 5 | 1 | 30 | 4 |
| SUB10 | 63 | 3 | 2 | 17 | 9 |
| SUB11 | 95 | 2 | 1 | 15 | 0 |
| SUB12 | 92 | 8 | 0 | 19 | 0 |
| SUB13 | 91 | 5 | 4 | 30 | 0 |
| SUB14 | 81 | 4 | 5 | 9 | 3 |
| SUB15 | 150 | 7 | 0 | 46 | 6 |
| SUB16 | 51 | 5 | 0 | 12 | 0 |
| SUB17 | 79 | 0 | 0 | 13 | 0 |
| SUB18 | 102 | 3 | 0 | 37 | 7 |
| SUB19 | 85 | 2 | 7 | 30 | 6 |
| SUB20 | 78 | 4 | 0 | 39 | 16 |
| SUB21 | 113 | 5 | 3 | 46 | 7 |
| SUB22 | 118 | 2 | 4 | 30 | 9 |
| SUB23 | 95 | 3 | 0 | 25 | 0 |
| SUB24 | 57 | 6 | 2 | 12 | 0 |
| SUB25 | 92 | 7 | 1 | 18 | 6 |
| SUB26 | 48 | 0 | 0 | 8 | 4 |
| SUB27 | 70 | 5 | 0 | 21 | 0 |
| SUB28 | 111 | 6 | 3 | 27 | 11 |
| SUB29 | 87 | 7 | 2 | 27 | 9 |
| SUB30 | 85 | 1 | 1 | 24 | 6 |
| SUB31 | 103 | 7 | 10 | 24 | 2 |
| SUB32 | 141 | 8 | 0 | 27 | 0 |
| SUB33 | 71 | 7 | 1 | 25 | 1 |
| SUB34 | 86 | 8 | 2 | 27 | 5 |
| SUB35 | 127 | 9 | 7 | 14 | 12 |
| SUB36 | 117 | 13 | 6 | 45 | 4 |
| SUB37 | 78 | 1 | 1 | 19 | 1 |
| SUB38 | 98 | 4 | 5 | 27 | 11 |
| Total | 3317 | 181 | 92 | 902 | 178 |

